# PeptiLine: an interactive platform for customizable functional peptidomic analysis

**DOI:** 10.1101/2025.10.23.684139

**Authors:** Russell Kuhfeld, Søren D-H. Nielsen, David C. Dallas

## Abstract

The PeptiLine peptidomics pipeline is a comprehensive software platform designed to transform peptidomic mass spectrometry output data into interactive, intuitive visualizations and statistical summaries. By integrating peptide-to-protein mapping, quantitative comparisons, bioactive annotations and descriptive and inferential analyses into a single user-friendly interface, PeptiLine addresses the bottleneck of timely and complex data exploration and visualization in bioactive peptide research. It enables visualization of peptidomic data through a series of modules allowing the user to create peptide sequence heatmaps, correlation plots and a series of categorical bar plots and pie plots to explore the absolute and relative bioactivity, protein origin, summed abundance and total peptide count. Here, we provide example uses of our Data Transformation, Descriptive Analysis and Heatmap Visualization tools to pinpoint functional peptide hotspots across protein sequences and compare samples across different protein variants. PeptiLine is freely available at https://mbpdb.nws.oregonstate.edu/peptiline/ and on GitHub (https://github.com/Kuhfeldrf/peptiline/) under an MIT license. The software runs on Windows, macOS, and Linux systems with Python ≥3.10. Support and bug reports.

**Author summary:** Bioactive peptides in food have health benefits ranging from antimicrobial activity to blood pressure regulation. We use mass spectrometry to identify these peptides, but analyzing the resulting datasets is challenging. The data comes out as massive spreadsheets that are difficult to interpret, and existing tools either require advanced programming skills, lack key analytical features, or are no longer maintained.

We built PeptiLine to address these problems. It’s a user-friendly platform that transforms peptidomic data into interactive visualizations and statistical summaries. The key challenge we solved was handling peptides with multiple overlapping functions, for instance, a peptide that’s both antimicrobial and antihypertensive. Most tools would double-count such peptides, inflating the totals. We figured out how to accurately display both individual functional categories and true abundance totals without this double-counting bias.

We demonstrated PeptiLine’s capabilities by analyzing bitter peptides in aged Cheddar cheese, successfully identifying specific protein regions responsible for bitterness and novel insight into Cheddars bioactive potential. The software is freely available as both a web application and for local installation, requiring no programming expertise.

## Introduction

Proteins from an array of different species, including dairy, plant, marine and animal proteins, are sources of encrypted peptides with diverse functions, including antimicrobial, antihypertensive, antioxidant, immunomodulatory, opioid and numerous others [1–5].

Researchers annotate MS/MS-based peptide data through functional peptide prediction models and databases with known functional peptides. Many different curated databases exist containing peptides and their corresponding known bioactivities, such as the Milk Bioactive Peptide Database (MBPDB) for milk peptides [3–4] or BioPepDB [5] for functional food peptides. In addition, over 200 different prediction models ranging over 17 bioactive functions have been published [6] with anticancer, antimicrobial and antiviral being the most prevalent bioactive functions.

While these tools help identify possible bioactive peptides in samples, they often lack integrated analytical tools for processing and visualizing experimental peptidomic datasets. To address this limitation, we developed PeptiLine, a comprehensive computational platform that integrates peptide annotation with visualization and statistical analysis tools. PeptiLine is hosted within the MBPDB but is designed to be broadly applicable across all peptidomic research domains.

### Current limitations in peptidomic analysis

Peptidomic data analysis is limited by several challenges. The variety of file formats produced by different mass spectrometry tools creates compatibility barriers between software applications [7]. Peptidomic data analysis requirements differ from those of proteomic data and therefore require specialized tools [8]. Many existing applications are difficult to install, require advanced computational expertise [9] and face continuity issues such as link rot and discontinued web servers, exemplified by the loss of platforms like Peptimetric [10].

### Gaps in current visualization tools

While existing peptidomic visualization platforms such as Peptimetric, Peptigram and PepMapViz have advanced the field [10–12], important limitations remain in their ability to translate large-scale data into biological insights. Peptimetric provides statistical analysis with built-in normalization but lacks functional data exploration capabilities and dedicated protein context mapping; moreover, its web application is inactive, and local installation requires technical expertise. Peptigram and PepMapViz generate useful sequence heatmaps but offer limited detail and customization, restricting their utility for nuanced peptide to sequence mapping.

A further challenge arises as bioactive peptides often have multiple, overlapping functions, which can lead to “double counting” when abundances are summed across functions. In datasets where a single peptide is mapped to multiple functions, totals can differ depending on whether abundances are summed by peptide or by function. Conventional charts such as pie plots or relative abundance bar plots often obscure this overlap, introducing bias into interpretation. Critically, no existing visualization tools are specifically designed to represent and explore overlapping functional data transparently.

High-throughput peptidomic experiments generate large, complex datasets that are difficult to interpret [13]. Current approaches also lack cost-effective methods for predicting, profiling and screening bioactive protein hydrolysates [14]. Together, these gaps highlight the pressing need for new visualization strategies tailored to functional peptidomic data.

### PeptiLine solution

To overcome the limitations of current visualization tools, we created PeptiLine, a comprehensive platform designed specifically for functional peptidomic analysis. PeptiLine’s modular architecture addresses these challenges through three integrated components for data transformation, descriptive analysis and sequence visualization (Fig 1). Through a representative case study, we demonstrate PeptiLine’s ability to address key challenges in visualizing and interpreting complex peptide datasets.

**Fig 1.**
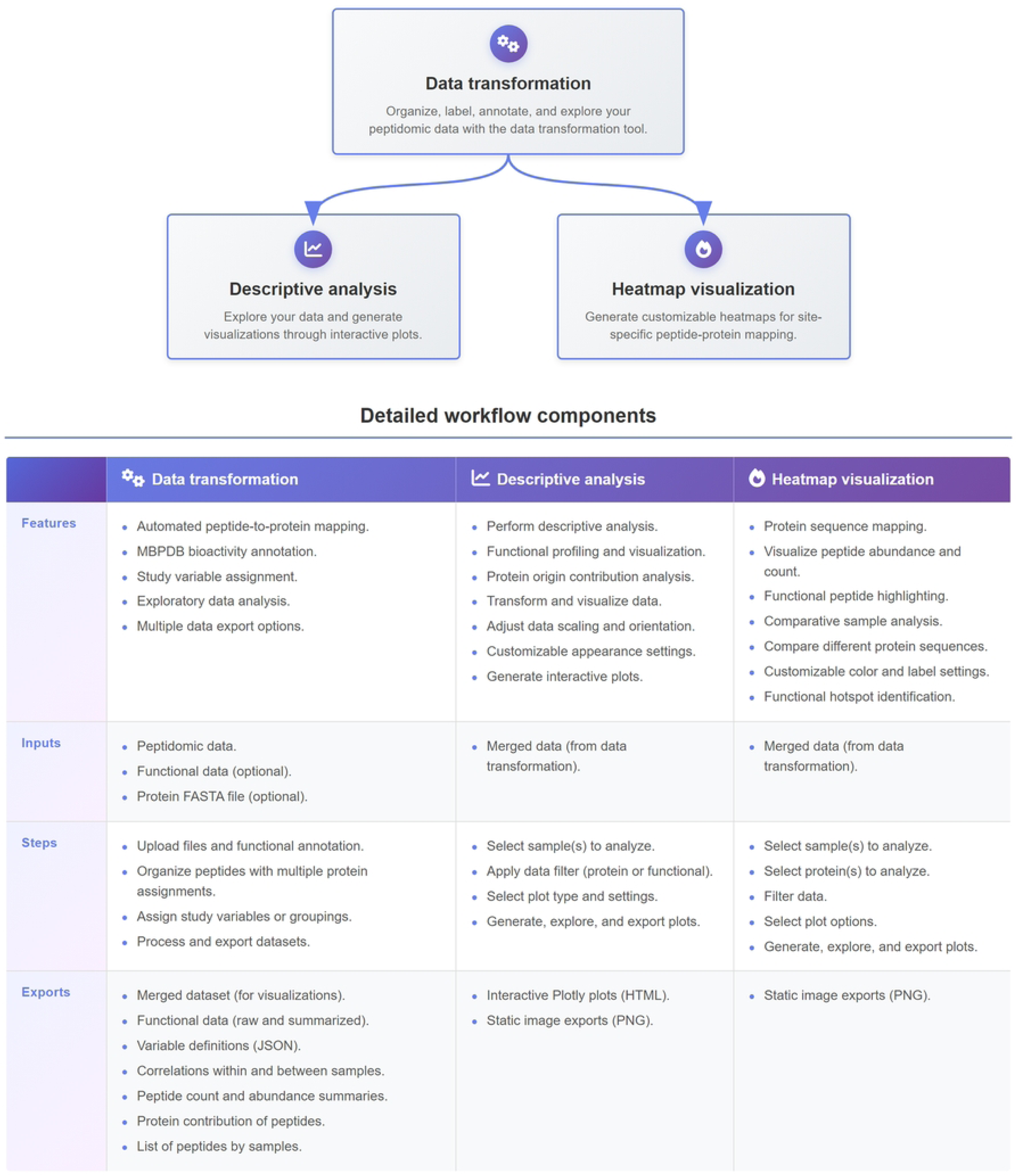
Comprehensive workflow for peptidomic data analysis using the PeptiLine platform.

### Case study

To demonstrate PeptiLine’s practical utility, we applied it to a publicly available peptide dataset (https://github.com/Kuhfeldrf/Bitterness-in-Aged-Cheddar-Cheese) [15]. PeptiLine streamlined core tasks such as variable assignment and comparison, annotation and abundance profiling while also enabling functional analysis of bioactive properties, showing its value for both general peptidomic workflows and targeted functional studies.

### Design and Implementation

#### Design

PeptiLine is designed to be easy-to-use and flexible, implemented as an integrated pipeline deployed as interactive Jupyter Notebook applications. By combining code, results and explanation in one place, it allows for easy customization [16] that is often lacking from other peptidomic workflows.

The platform is developed in Python (≥3.10) with Jupyter Notebooks 7.4.5 and uses established scientific libraries such as Pandas, NumPy, Matplotlib, Seaborn, Plotly and Scikit-learn. IPyWidgets are incorporated as the user interface of notebooks giving a web-application experience.

PeptiLine’s modular architecture consists of three integrated components (Fig 1): (i) Data Transformation for organizing and annotating peptidomic data with automated handling of multi-protein peptide assignments and MBPDB bioactivity annotation; (ii) Descriptive Analysis for exploratory data analysis through interactive visualizations; and (iii) Heatmap Visualization for creating visual maps of peptide density and positioning along the linear structure of parent proteins with customization options.

### Implementation

Within the MBPDB web application, Jupyter Notebooks are deployed via Voilà which renders Jupyter Notebook into an interactive web application, eliminating the need for local installation. For local implementation, the GitHub repository (https://github.com/Kuhfeldrf/peptiline/) contains installation instructions (<u>README.peptidline.md</u>), dependencies (<u>notebook_requirements.txt</u>) and the three module notebooks (<u>data_transformation.ipynb</u>, <u>heatmap_visualization.ipynb</u>, <u>data_analysis.ipynb</u>) with a supporting file (<u>_settings.py</u>) and folders such as the UniProt API utilities (<u>utils</u>) and case study data (<u>examples</u>). Local installation removes direct MBPDB querying capability.

### System requirements

PeptiLine runs on any modern computer supporting Jupyter Notebooks (Windows, macOS, Linux). The web-based version requires only a modern web browser and internet connection. For local deployment, users need Python ≥3.10, Jupyter Notebook, and the dependencies listed in notebook_requirements.txt.

## Results

### Description of modules and case study example

#### Data Transformation

The Data Transformation module standardizes peptidomic outputs from multiple proteomics platforms (Proteome Discoverer, MaxQuant, Skyline, FragPipe, etc.) by accepting common file formats (CSV, TXT, TSV, XLSX) with flexible delimiters and automatically mapping software-specific column names to a unified format (Table 1, Fig 2A). Each peptide is assigned a unique identifier that combines its sequence with any modifications, preventing redundancy and enabling consistent tracking across datasets.

**Fig 2.**
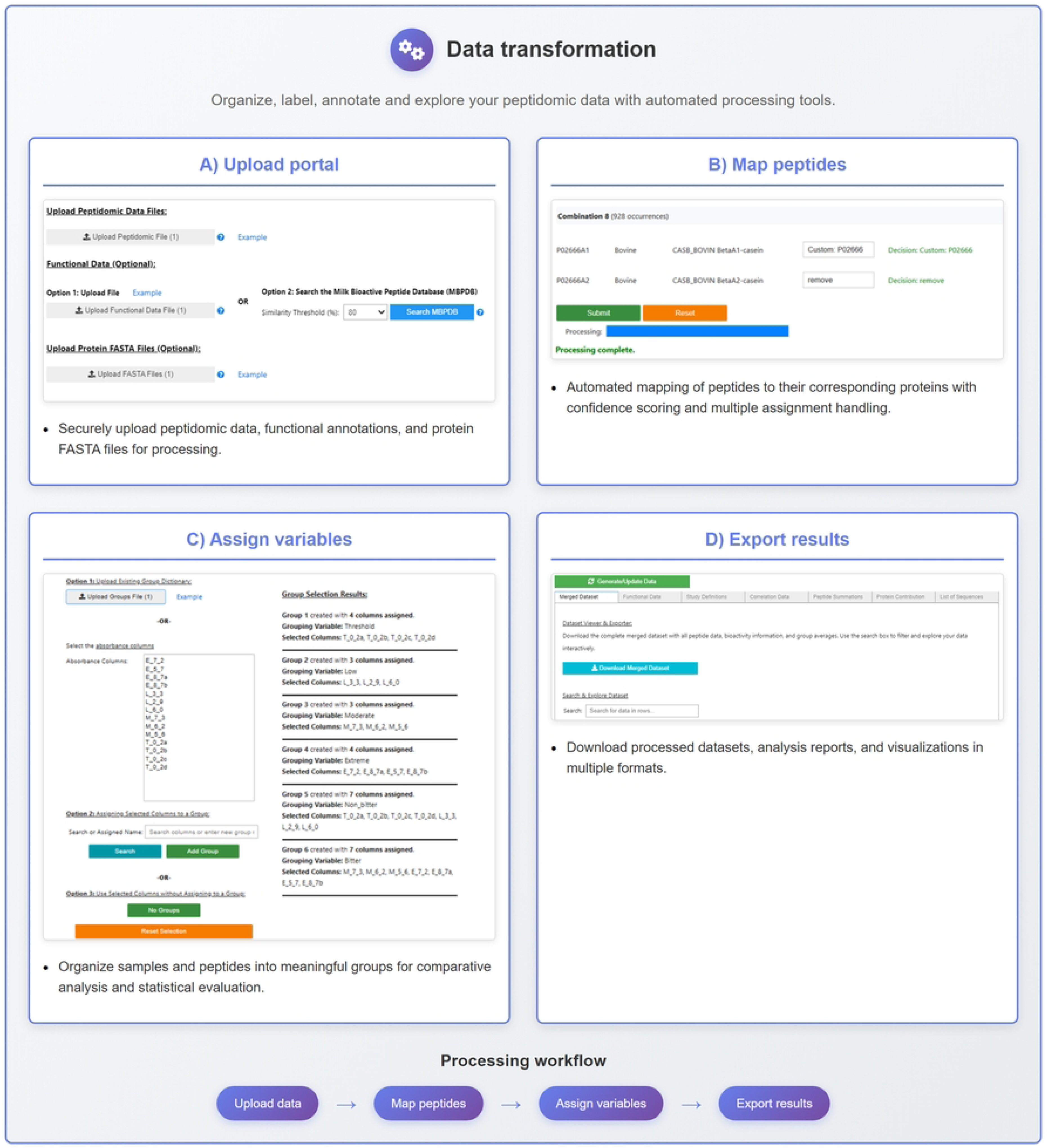
Data transformation module core workflow.

**Table 1.**
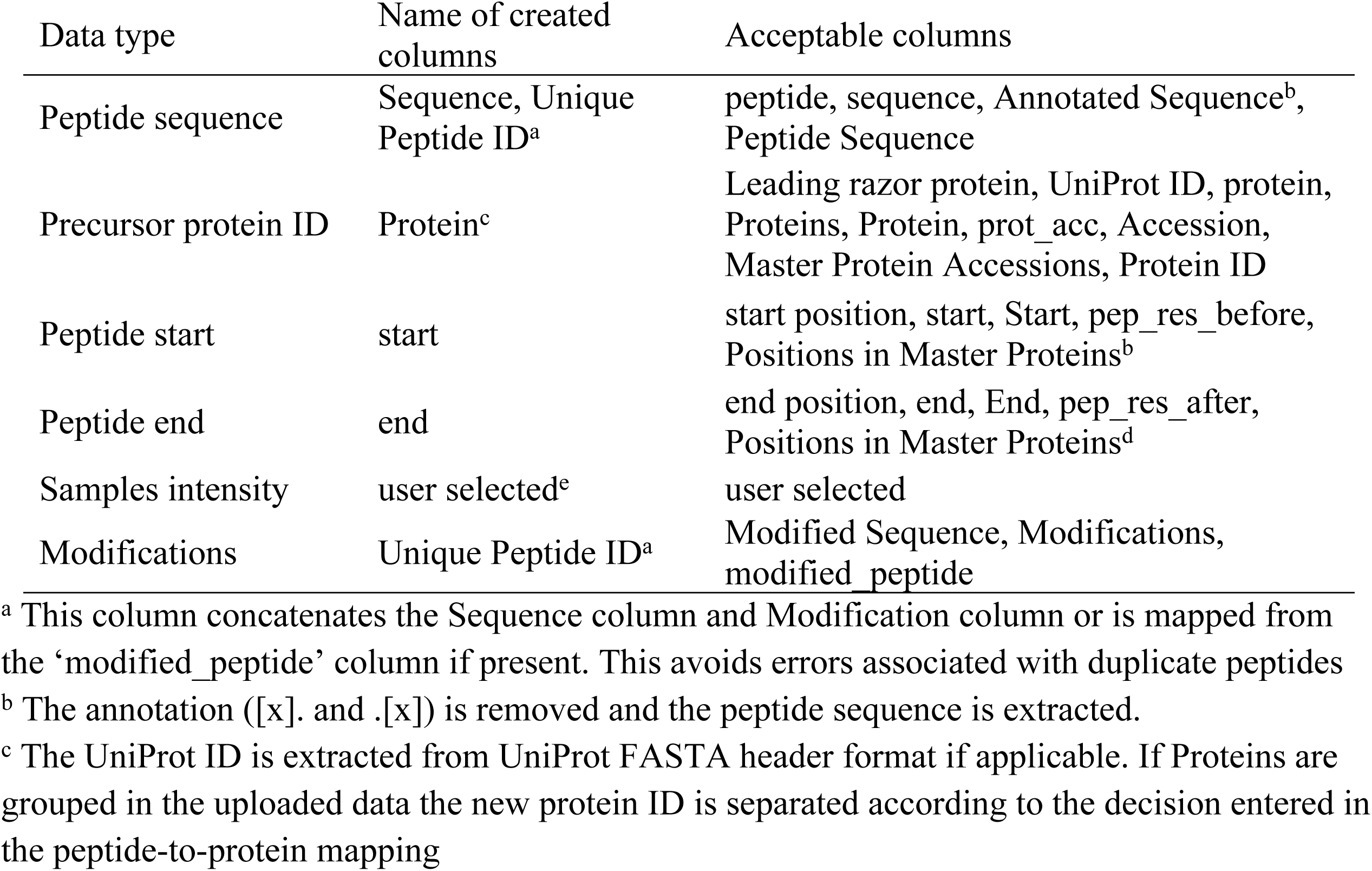
Supported column mapping for peptidomic inputs from a wide range of software applications.

Functional annotation is achieved either by integrating external datasets with required columns and column mapping detailed in Table 2 or by querying the MBPDB, where peptides are annotated with known biological functions based on exact matches or user-defined homology thresholds. To illustrate the module’s functionality, we applied it to the case study’s <u>peptidomic</u> <u>data</u>. The dataset was searched against the MBPDB to identify and annotate bioactive peptides.

**Table 2.**
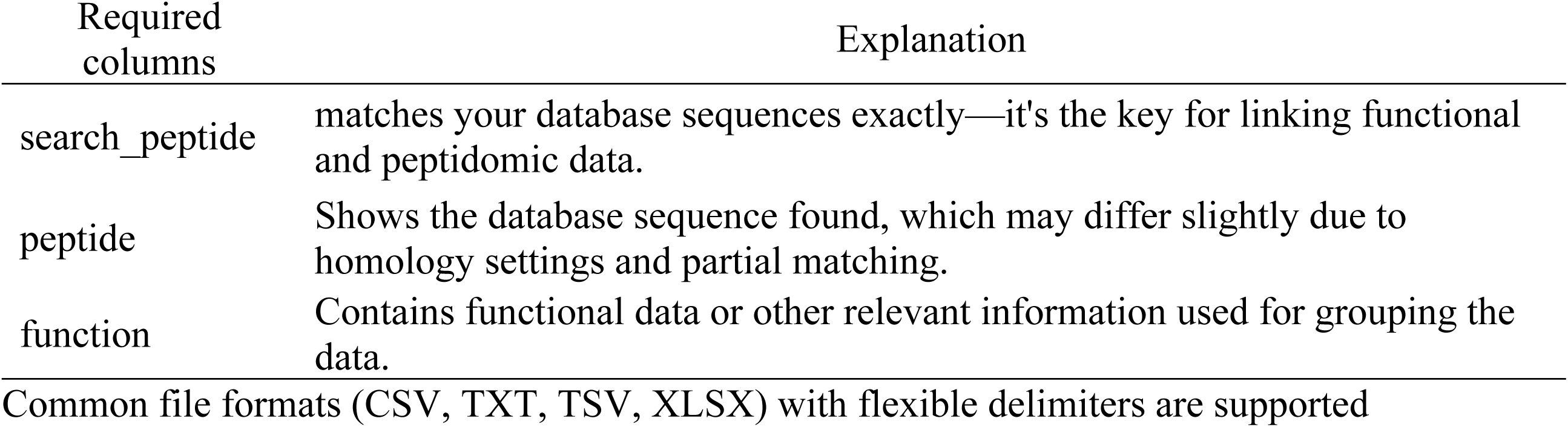
Required Columns for Functional Database Integration.

To address peptides mapping to multiple proteins, the study’s <u>protein library</u> was first uploaded and then the module’s peptide-to-protein mapping tool (Fig 2B) was used to split, retain or remove ambiguous mappings. This functionality is particularly critical for β-casein analysis in the case study, where variants A1 and A2 differ by a single amino acid substitution at position 67 (histidine in A1, proline in A2). The peptide-to-protein mapping tool was used to organize shared peptides for both combined and variant-specific analyses. For instance, peptide NSLPQNIPPL, originally mapped by Proteome Discoverer to “P02666A1; P02666A2 at positions [68–77],” was consolidated to “P02666 [68–77]” for aggregate β-casein analysis (Fig 2). This transformability ensures consistent peptide-to-protein assignments while maintaining flexibility to analyze both combined and individual variant contributions.

Users assign study variables through an interactive interface or JSON dictionaries that map experimental groups to abundance columns, automatically calculating averaged abundance values across replicates while retaining raw data for audit trails. In the case study, bitterness groups (Threshold, Low, Moderate, Extreme) and broader categories (Non_bitter = Threshold + Low; Bitter = Moderate + Extreme) were mapped to abundance columns using the variable assignment tool (Fig 2C) following the case study’s <u>sample overview table</u>. Categorical integration enabled grouped visualizations and statistical comparisons while maintaining data traceability. Multiple grouping schemes can be applied simultaneously, enabling rapid exploration of functional peptide distributions across experimental conditions.

The module generates comprehensive exports including a merged dataset that integrates peptide identifications, abundances, functional annotations and study variables. This unified dataset serves as the primary file for subsequent analyses, ensuring that downstream visualization and statistical comparisons are based on complete and consistently organized data. Additional outputs include functional summaries, group definitions, correlation analyses, protein-level abundance distributions and peptide sequence lists (Fig 2D, Tables S1-S10).

### Descriptive Analysis

The Descriptive Analysis module enables comprehensive exploration of peptide abundance and functional profiles across samples. Users begin by uploading their processed data (merged dataset; Table S2) (Fig 3A) and selecting sample groups for comparison (Fig 3B). Prior to visualization, several optional data filtration options are available (Fig 3C), including restricting output to chosen proteins, bioactive functions or both. An additional option allows peptides to be categorized as functional or non-functional based on annotated bioactivities. The Descriptive Analysis module provides flexible visualization options including grouped/stacked bar charts, pie charts and correlation scatter plots. Data can be displayed as absolute values or relative percentages for peptide count or abundance, oriented by sample, protein or function. All visualizations include correlation assessments (Pearson or Spearman, with optional log transformation) and generate publication-ready interactive or static figures (Fig 3C).

**Fig 3.**
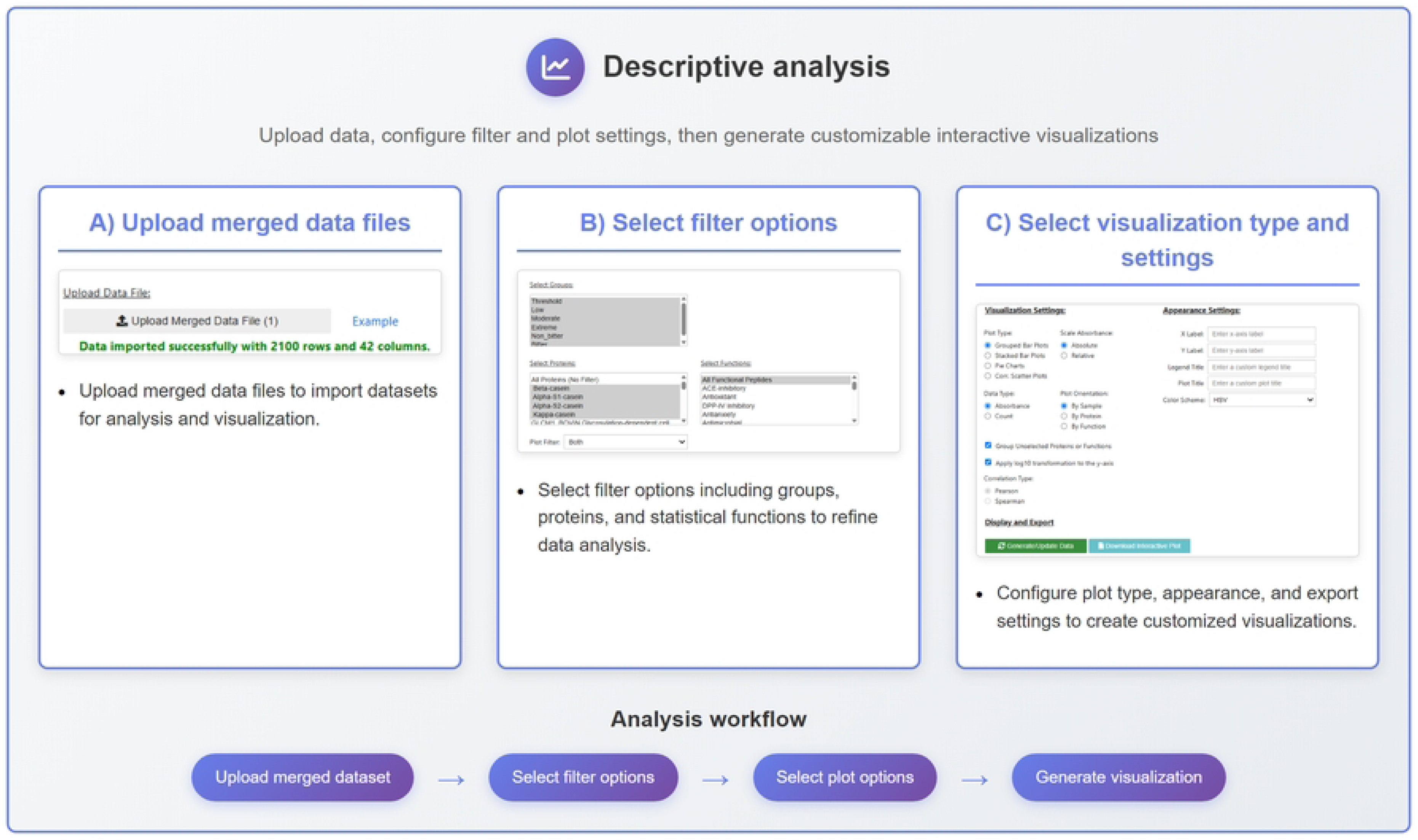
Descriptive analysis module core workflow.

### Case Study: Correlative Analysis

Sample relationships are explored through interactive scatter plots, scatter plot matrices (SPLOMs) (Figs 4A and S1) and correlation tables (Table S6). This revealed that the Threshold group showed the most distinct peptidomic profile, with weaker correlations to other bitterness levels (r < 0.718). In contrast, Low, Moderate, and Extreme groups were strongly correlated (r > 0.824), indicating similar peptide release patterns among bitter samples. This suggests that bitterness perception involves a fundamental shift in peptide composition at the Threshold-to-Low transition, with higher bitterness levels representing much smaller differences in peptide profiles.

**Fig 4.**
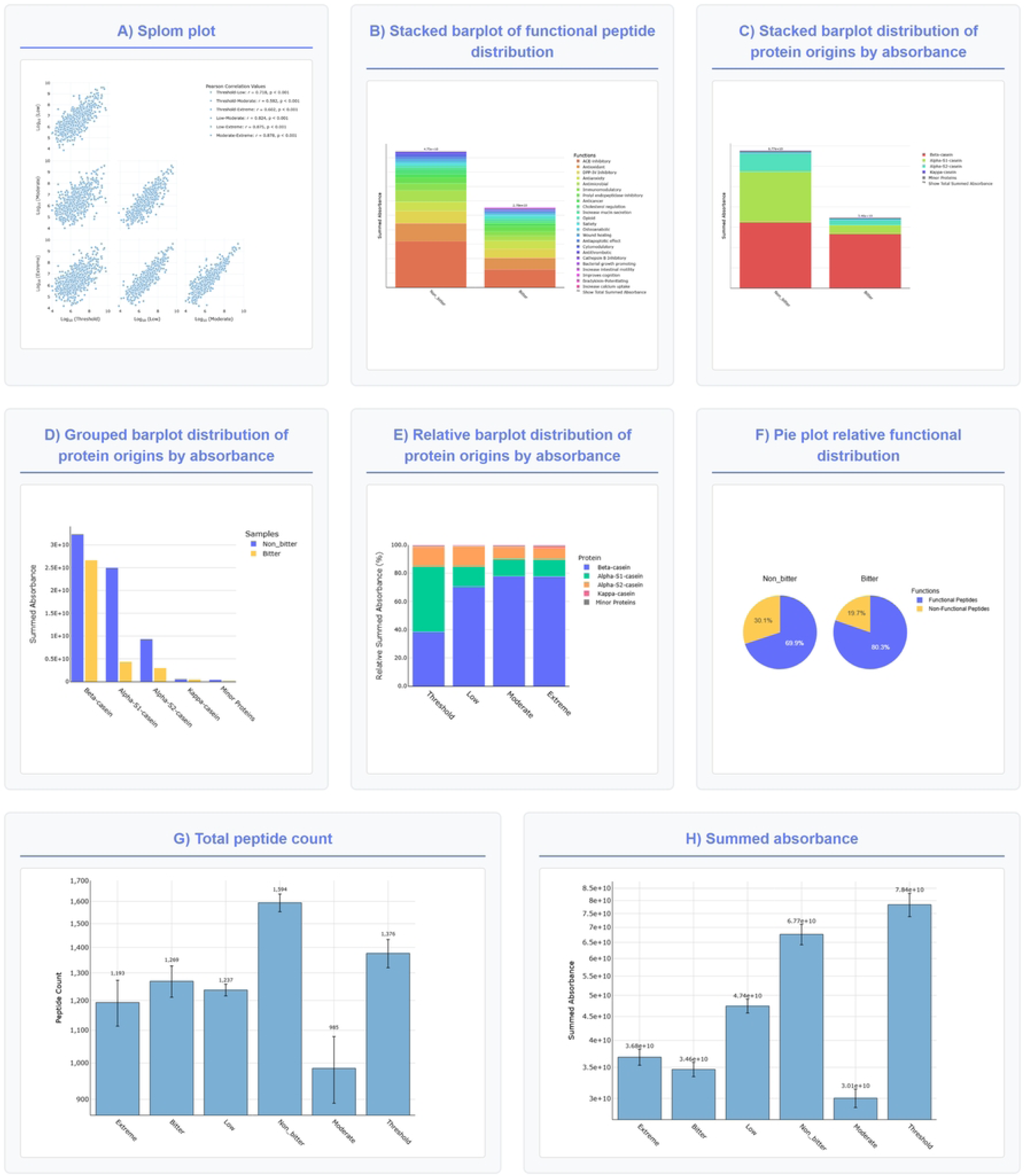
Example of descriptive visualization generated from the data analysis module.

### Case Study: Functional Analysis

Leveraging the integrated functional annotations, this module categorizes peptides by their known functions. The primary visualization is an interactive stacked bar plot (Figs 4B and S2; Table S4) that displays functional peptide distributions across samples, enabling direct comparison of bioactive profiles between experimental groups.

Analysis of the cheese dataset revealed distinct bioactive profiles between bitter and non-bitter samples across 21 functional categories (Figs 4B and S2; Table S4). ACE-inhibitory peptides dominated both groups but showed 37% lower abundance in bitter samples despite similar peptide counts (350 vs. 332 peptides). Several functions exhibited different patterns: antioxidant activity was similar across bitter and non-bitter samples (+4%), antimicrobial peptides were lower in the bitter group (−33%), while DPP-IV inhibitory activity was higher in the bitter group (+20%). Overall, total bioactive peptide abundance was 41% lower in bitter samples, suggesting that proteolytic degradation during cheese aging selectively enhances certain bioactivities while reducing others and overall functional peptide concentrations. These patterns demonstrate how visualization tools can reveal functional consequences of food processing conditions on bioactive peptide profiles.

### Case Study: Protein Origin Contribution Analysis

The key sources of peptides in this case study are explored through the scaled bar plots (Figs 4C, 4D, 4E and S3, Table S9) that simultaneously display absolute abundance and relative contribution percentages. This visualization allows unbiased comparison of which proteins contribute with most peptides within samples and across experimental conditions. In the case study, this visualization showed that β-casein-derived peptides had similar measured abundances between non-bitter and bitter samples (3.24 × 10¹⁰ vs. 2.67 × 10¹⁰). In contrast, the scaled visualization shows that β-casein’s relative contribution amongst other detected peptides was substantially higher in bitter samples (76.86%) than in non-bitter samples (47.79%) (Figs 4C and S3, Table S9). This dual visualization approach demonstrates how examining both relative proportions and measured abundances together provides a more complete understanding of the underlying degradation mechanisms.

### Case Study: Total Peptide Counts and Abundance Summaries

The count of unique peptides and summed abundances across sample groups are visualized through bar charts with error bars representing replicate variability (Figs 4G, 4H, S4, S5, Table S8). The analysis revealed that non-bitter samples contained more unique peptides (1,594) and higher total abundance (6.77 × 10¹⁰) than bitter samples (1,269 peptides, 3.47 × 10¹⁰ abundance), suggesting that peptide diversity and abundance decrease during bitterness development in aged cheese.

### Sequence Visualization

The Sequence Visualization module provides a novel approach to simultaneously compare peptide abundance and site-specific peptide counts by integrating both metrics into a single heatmap representation. The processed data (merged dataset; Table S2) was uploaded to the module along (Fig 5A) with the protein sequences via FASTA files or integrated UniProt queries, then the user selects the target proteins and experimental groups with customizable labeling and sample ordering options (Fig 5B). Visualization parameters include peptide filtering (all peptides, functional peptides, or specific functions) and discrete color options to best illustrate data trends (Fig 5C).

**Fig 5.**
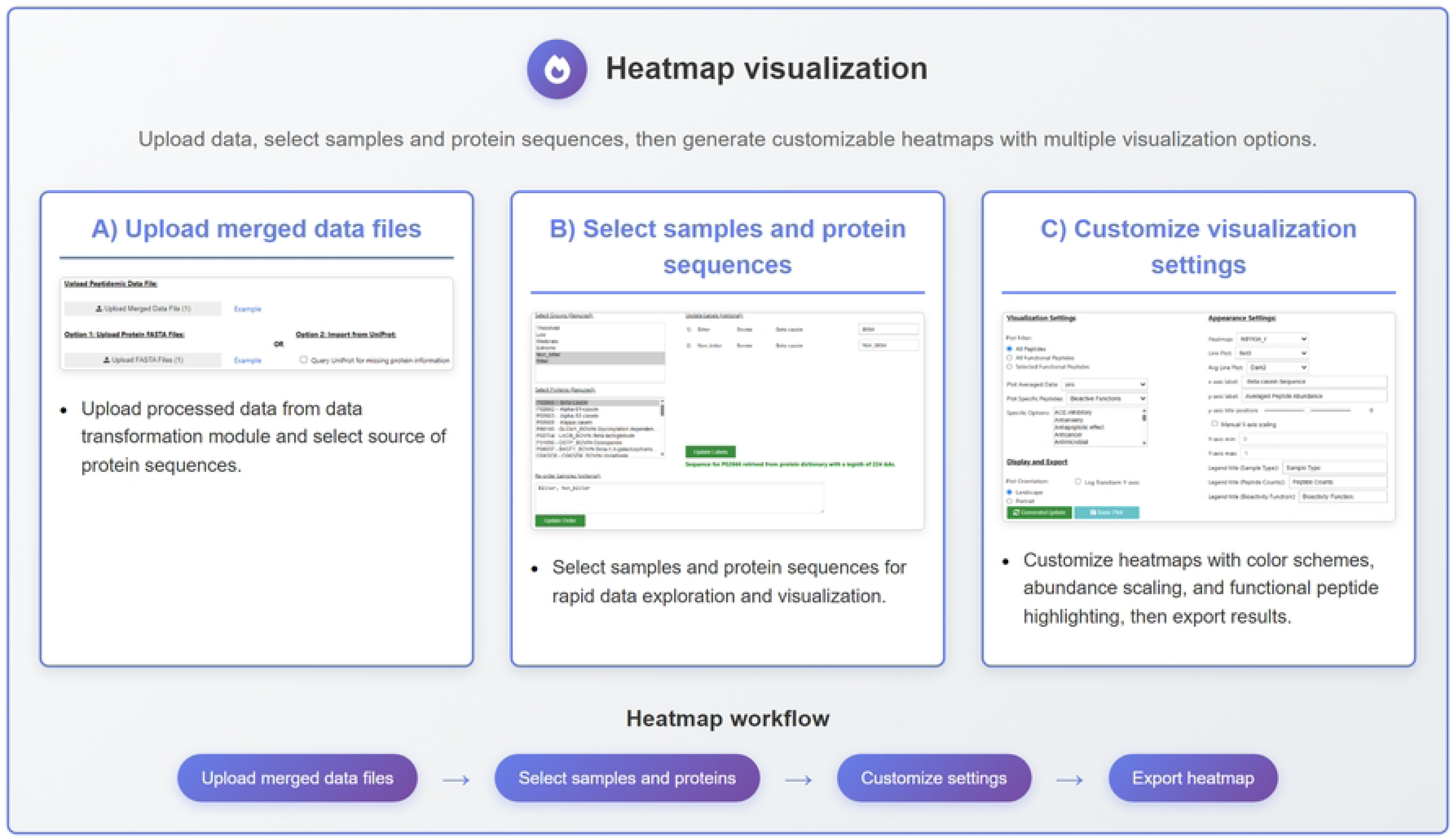
Heatmap visualization module core workflow.

The sequence heatmaps (Fig 6A) are generated through a multi-step process: the filtered merged dataset (Fig 6B) is first transformed to calculate positional count and absorbance values (Fig 6C), then visualized to display amino acid position on the x-axis and averaged peptide abundance on the y-axis, with color intensity indicating peptide count frequency at each sequence position. This enabling comparison between experimental groups and bioactive functions in context of their position within a protein sequence. This integration reveals unique patterns in proteolysis between experimental conditionals, functionally important protein regions and bioactive peptide clustering patterns.

**Fig 6.**
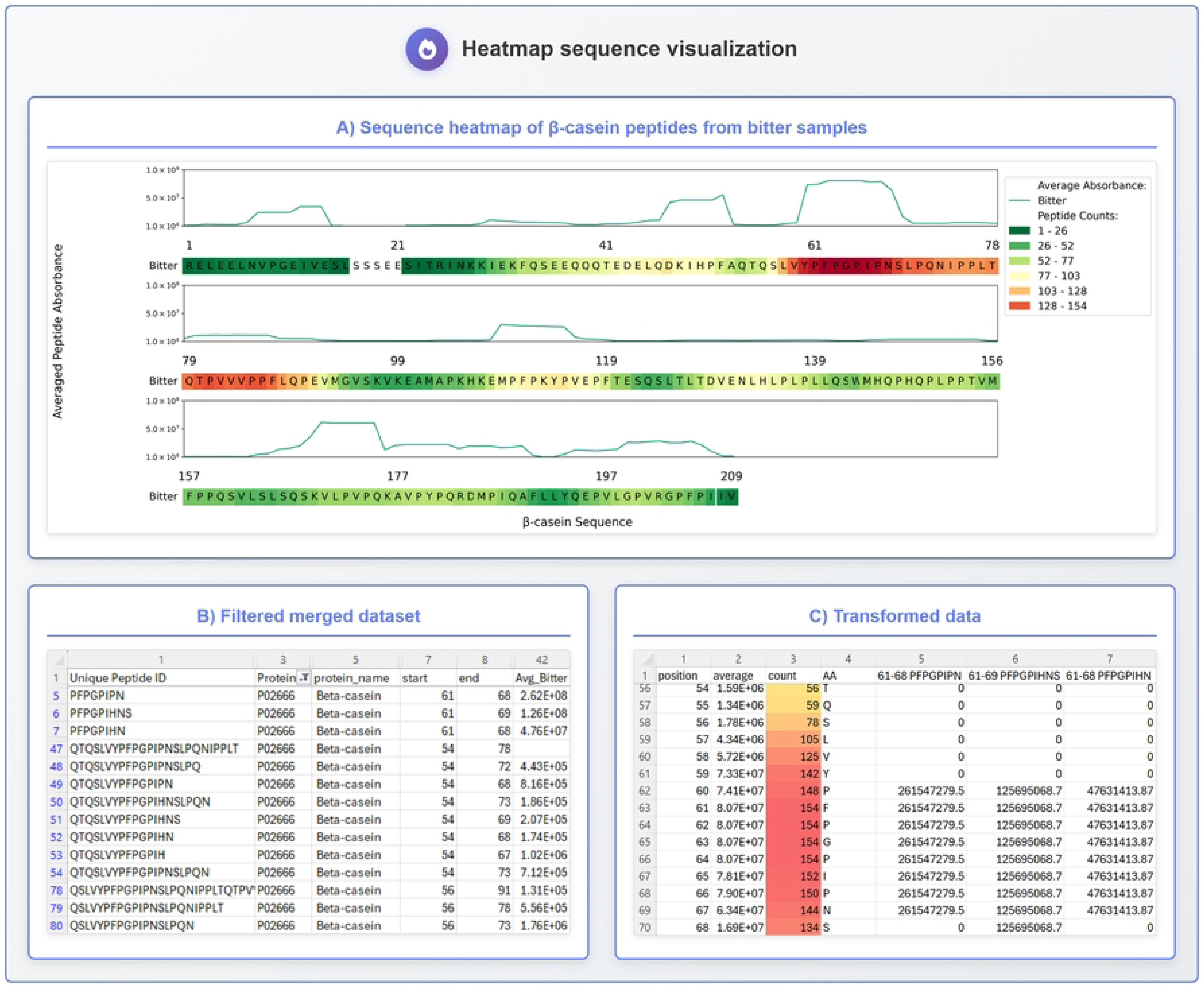
Heatmap visualization sequence heatmap example.

The utility of this visualization was demonstrated by analysis of bitter β-casein-derived peptides in aged Cheddar cheese, a major contributor of both bitter peptides [17] and bioactive peptides [3]. The heatmap revealed amino acids 60–67 of β-casein as having relatively high peptide abundance (Fig 4 A). This region emerged as a “hotspot” of bitterness, with abundance overlays showing that despite similar peptide numbers in the 60–67 region, the average abundance of bitter cheeses was notably higher than non-bitter cheeses. Additionally, region 60– 67 contributed the most peptides across the entire β-casein sequence.

Figure 7A shows the overall distribution of all β-casein-derived peptides with A1 and A2 variants aggregated, showing the 60–67 regions as a distinct source of elevated peptide abundance in bitter samples compared to non-bitter ones. PeptiLine’s custom FASTA file capability enabled analysis of peptides mapped exclusively to β-casein variants A1 and A2 (Fig 7B), addressing limitations in traditional mapping applications. As UniProt protein ID ‘P02666’ maps to the A2 variant (proline at position 67), custom uploads were essential for analyzing the A1 variant (histidine substitution) and for removing signal sequences which is the norm in bitter peptide research nomenclature. The variant comparison revealed that β-casein A1 contributed more bitter peptides than A2, consistent with previous findings [17].

**Fig 7.**
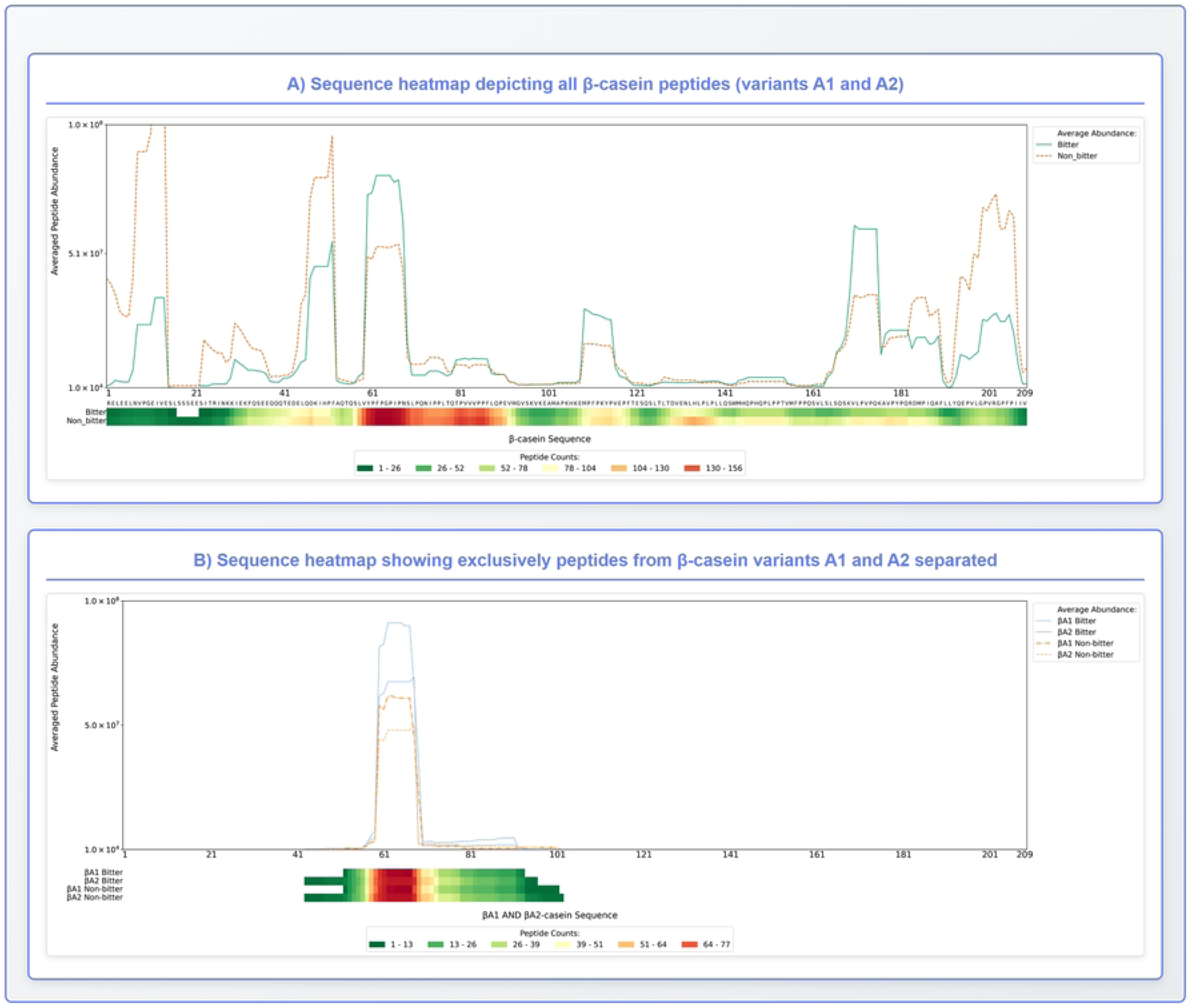
Heatmap visualization of peptide abundance and density across β-casein sequence in bitter and non-bitter samples. A) All β-casein peptides (A1 and A2 variants). B) Variant-specific peptides.

The module incorporates comprehensive error handling, supports custom sequence uploads and provides interactive guidance to ensure accessibility across varying computational expertise levels offering an improvement over existing tool like Peptigram.

### Comparison with other tools

#### Sequence Heatmap

The sequence heatmap generated by PeptiLine (Fig 7B) provides enhanced visualization capabilities compared to Peptigram’s approach (Fig S6). While Peptigram combines peptide intensity and count into a single compact visualization, this simplicity reduces information depth and sample comparison detail. More critically, PeptiLine prevents erroneous peptide mapping that occurs with other tools. For example, Peptigram incorrectly mapped the case studies data set showing β-casein A1 peptides around position 20 (Fig S6), when the actual sequence coverage spans positions 43-102 (Fig 7B).

PeptiLine addresses these limitations through independent display of peptide count and abundance with overlay capabilities, enabling detailed sample comparisons across full protein sequences. Technical advantages include robust handling of missing data (avoiding Peptigram’s bias-introducing zero imputation), support for custom protein sequences/variants not available in standard databases and advanced customization options.

The automated system represents a substantial improvement over manual Excel-based workflows [3,4,15,17], rapidly generating fully annotated heatmaps with flexible plot orientation (portrait or landscape) and the ability to highlight bioactive peptide regions. These enhancements collectively streamline sequence-based analyses while improving accuracy and interpretability of peptidomic data. Notably, PeptiLine is the first sequence heatmap visualization tool specifically designed with integrated features focused on functional peptide annotations, allowing researchers to map bioactive properties directly onto protein sequences.

### Functional Analysis

A key challenge in functional peptide analysis is that bioactive peptides often have multiple overlapping functions, leading to “double counting” when abundances are summed across functions. Furthermore, total functional peptide abundance or count per sample is often calculated differently if summed across peptides (i.e., sum of all bioactive sequences’ abundance) vs. if summed across function (e.g., sum of antioxidant sequences’ abundance), creating challenges in how to visualize the data. Conventional charts such as pie plots [18] (Fig 4F) or relative bar charts (Fig 4E) often obscure this overlap, introducing bias into interpretation, while no existing tools are designed to represent overlapping functional data transparently.

PeptiLine addresses these visualization challenges through interactive scaled abundance plots that simultaneously display both relative and absolute abundances, enabling accurate sample-to-sample and within-sample comparisons. The key innovation is scaling the total height of each bar based on unique peptide summation (Totals row in Table S4, Total Summed Absorbance in Fig S2) while depicting the relative abundance of functions within each sample based on functional summation (Rows 3-25 in Table S4, items in legend in Fig S2). This visualization approach creates a Y-axis scaled figure that allows accurate sample-to-sample comparison while showing the qualitative relative contribution of each function within samples or across samples given the scaling according to absolute abundance or count.

Unlike manual approaches requiring researchers to create separate visualizations in Excel, R or Python to determine optimal data representation, PeptiLine enables rapid iteration through comprehensive visualization options (grouped vs. stacked bar plots, pie charts by sample or function, count vs. abundance metrics) within a single interface. This functionality eliminates the time-consuming manual plot creation process and prevents suboptimal visualization choices that can compromise biological interpretation.

### Relative Protein Contribution Analysis

The scaled abundance plots such as those from [19] and generated by PeptiLine (4C, and S3) offer distinct advantages in visualizing the amount of a source protein relative to other proteins within a sample and between samples in sample-to-sample comparisons. Traditional approaches commonly used in peptidomic literature, such as pie charts or relative abundance bar plots for source show only the proportion of each protein within a sample without accounting for total peptide abundance differences between samples [20]. While PeptiLine provides pie charts and relative abundance bar plots (Figs 4E and 4F) as standard visualization options, the scaled bar plots offer unique advantages for comprehensive data interpretation.

A key distinguishing feature of PeptiLine is its use of the peptide data table exported from peptidomic processing software rather than traditional protein abundance tables. This approach enables users to first address ambiguous mapping of peptides to multiple proteins, providing researchers the flexibility to access aggregated contributions from all protein variants.

Through the peptide-to-protein mapping tool and data visualization, one can easily assess the contribution of all protein variants either combined (e.g., all β-casein variants together) or separated (e.g., β-casein A1 vs. β-casein A2). This peptidomic data approach provides a more flexible approach to viewing how different protein forms contribute to the overall peptide profile.

The scaled abundance visualization particularly excels at revealing differential protein contributions across experimental conditions while maintaining the ability to compare both absolute and relative abundances simultaneously. This dual perspective is critical for identifying proteins that may appear similar in relative abundance, but differ significantly in absolute contribution, providing researchers with a more complete understanding of the protein source of bioactive peptides.

### Limitations of Count-Only Analysis

Approaches that rely solely on peptide counts for protein source or functional assessment are commonly employed in peptidomic literature but introduce significant analytical limitations [21–24]. These traditional count-based metrics disproportionately emphasize low-abundance peptides, creating misleading representations where peptide presence is weighted equally regardless of abundance.

This limitation is clearly demonstrated in our case study data, where count-only analysis would significantly underestimate α_S1_-casein’s biological contribution. In the bitter samples α_S1_-casein represents only 12.51% of total peptide count, it accounts for 23.33% of relative abundance, indicating that although this protein generates fewer distinct peptides, in aggregate these peptides contribute substantially more to the overall peptide profile. Additionally, the dominant bioactivity ‘ACE-inhibitory’ showed 37% lower abundance in bitter samples despite similar peptide counts (350 vs. 332 peptides; a 5% difference) compared with the non-bitter samples. This discrepancy illustrates how count-only approaches can distort the takeaways from functional data by placing an equal contribution to peptides with changing levels of abundance across samples or experimental conditions.

Approaches based only on peptide counts ignore differences in their actual abundance levels. PeptiLine addresses these traditional limitations by providing both count and abundance metrics, enabling researchers to select appropriate data representations based on their specific research questions and biological context. The summed abundance approach effectively integrates both peptide count and individual peptide abundance, providing a comprehensive representation that captures both the diversity of peptides present and their quantitative contribution to the sample.

All results presented are fully reproducible using the test data (examples folder), and parameters available in the repositories readme file (https://github.com/Kuhfeldrf/peptiline/blob/main/README.md) and supplementary materials.

## Availability and future directions

### Availability

The PeptiLine pipeline is available on the MBPDB web platform (https://mbpdb.nws.oregonstate.edu/peptiline/) and the source code is deposited on GitHub (github.com/kuhfeldrf/peptiline).

### Future Directions

Future development will enhance PeptiLine’s accessibility, flexibility and interactivity. Planned improvements include flexible data import with dynamic column mapping to support diverse file formats (e.g., Spectronaut®, PEAKS®). We are also developing a compact heatmap module (<u>heatmap_compact.ipynb</u>) inspired by Peptigram for streamlined multi-sample comparisons.

PeptiLine’s modular Jupyter notebook architecture enables user customization and community contributions. Users can extend functionality by adding support for additional software formats and databases, with continued module development enhancing the platform’s analytical capabilities and adaptability.

### Software Deposition Statement

The complete source code, documentation, test data, and dependencies described in this manuscript have been archived and deposited as supplementary materials (Code_Repository.zip) to ensure permanent availability and reproducibility. The actively maintained version is available on GitHub (https://github.com/Kuhfeldrf/peptiline/).

## Supporting information

**S1 Fig. Scatter plot matrix showing peptide abundance correlations between bitterness groups.** Pairwise correlation analysis of peptide abundances across Threshold, Low, Moderate, and Extreme bitterness levels with Pearson correlation coefficients.

**S2 Fig. Stacked barplot of functional peptide distribution.** Distribution of 21 functional categories across bitter and non-bitter samples with scaled bar heights representing total summed absorbance.

**S3 Fig. Stacked barplot - distribution of protein origins by absorbance.** Contribution of protein sources (Beta-casein, Alpha-S1-casein, Alpha-S2-casein, Kappa-casein, Minor Proteins) to total peptide abundance in bitter and non-bitter samples.

**S4 Fig. Total peptide count.** Total unique peptide counts across bitterness groups with error bars representing replicate variability.

**S5 Fig. Total peptide abundance.** Summed peptide abundances across bitterness groups with error bars representing replicate variability.

**S6 Fig. Sequence heatmap generated by Peptigram of β-casein bitter peptides.** Comparison figure showing Peptigram’s visualization of β-casein variants A1 and A2 peptides, demonstrating mapping differences with PeptiLine’s approach.

**S1 Table. Data export specifications and file descriptions.** Complete listing of all data files generated by PeptiLine’s Data Transformation module with file names, formats, and contents.

**S2 Table. Merged dataset.** Complete dataset with peptide identifications, abundances, and functional annotations (merged_dataframe.csv).

**S3 Table. MBPDB results.** Bioactive peptide search results from MBPDB database or uploaded file (MBPDB_SEARCH.tsv).

**S4 Table. Summed functional data.** Aggregated functional data with abundance and count summaries (Processed_mbpdb_results.xlsx).

**S5 Table. Group definitions.** Sample grouping variables and experimental design definitions (categorical_variable_definitions.json).

**S6 Table. Sample-to-sample correlations.** Correlation coefficients between experimental groups (group_correlations_*.xlsx).

**S7 Table. Technical replicate correlations.** Correlation coefficients within experimental groups for technical replicates (correlation_analysis_*.xlsx).

**S8 Table. Summed peptide results.** Peptide abundance summaries (count and abundance) by experimental group (summed_peptide_results.xlsx).

**S9 Table. Protein analysis results.** Protein-level abundance distributions across groups (protein_absorbance_analysis.csv).

**S10 Table. List of sequences.** Extracted list of unique peptide sequences detected per experimental group (list_of_peptides_by_sequences.csv).

**S11 Code_Repository.zip.** Archived version 1.0 of the codebase downloaded form the GitHub repository on the date of submission.

Interactive HTML versions of Figs S1-S6 (generated with Plotly) are available for dynamic data exploration at: https://mbpdb.nws.oregonstate.edu/supplementals_tables_and_figures/

